# Insights into Diversity, Host-Range, and Temporal Stability of *Bacteroides* and *Phocaeicola* Prophages

**DOI:** 10.1101/2024.10.21.619336

**Authors:** Nejc Stopnisek, Stina Hedžet, Tomaž Accetto, Maja Rupnik

## Abstract

**Background:** Phages are critical components of the gut microbiome, influencing bacterial composition and function as predators, parasites, and modulators of bacterial phenotypes. Prophages, integrated forms of these phages, are prevalent in many bacterial genomes and play a role in bacterial adaptation and evolution. However, the diversity and stability of prophages within gut commensals, particularly in the genera *Bacteroides* and *Phocaeicola*, remain underexplored. This study aims to screen and characterize prophages in these genera, providing insights into their diversity, host range, and temporal dynamics in the human gut.

**Results:** Using a combination of three bioinformatic tools—Cenote-Taker 3, Vibrant, and PHASTER—we conducted a comprehensive analysis of prophages in *Bacteroides* and *Phocaeicola*. Cenote-Taker 3 identified the most diverse set of prophages, with significant overlaps observed between the tools. After clustering high-quality prophages, we identified 22 unique viral operational taxonomic units (vOTUs). Notably, comparisons between prophages identified in isolated bacterial genomes, metaviromes, and large public gut virome databases revealed a broader host range than initially observed in single isolates. Certain prophages were consistent across time points and individuals, suggesting temporal stability. All identified prophages belonged to the *Caudoviricetes* class and contained genes related to antibiotic resistance, toxin production, and metabolic processes.

**Conclusions:** The combined use of multiple prophage detection tools allowed for a more comprehensive assessment of prophage diversity in *Bacteroides* and *Phocaeicola*. The identified prophages were not only prevalent but also exhibited broad host ranges and temporal stability. The presence of antibiotic resistance and toxin genes suggests that these prophages may significantly influence bacterial community structure and function in the gut, with potential implications for human health. These findings highlight the importance of using diverse detection tools to accurately assess prophage diversity and dynamics.

## Introduction

Prophages dominate gut viromes and influence composition and function of gut bacteria as predators, parasites and modulators of bacteria phenotypes [1]. Gut virome of healthy individuals is stable, but individual specific and alterations in the gut virome impact human health [2].

Phages exhibit various life styles, from lytic to chronic infection, carrier state and lysogenic [3]. Prophages were found in 75% of more than 13.500 bacterial genomes and are particularly enriched in *Enterobacteriaceae* [4]. Dependent on environmental signals phages can initiate lytic or lysogenic cycle after infection. Recent studies demonstrated frequent signal-dependent prophage induction in various environments [5–7]. For instance, experiments in mouse models have shown that environmental stressors can trigger prophage induction, leading to increased bacterial lysis and horizontal gene transfer [8, 9]. Similarly, human gut studies have reported active prophage induction, contributing to microbial community dynamics and potentially impacting host health [10–12]. The connection of prophage induction and associated host diseases, such as IBD, Chron’s disease and Ulcerative colitis has also been suggested [13].

Free and integrated phages are in dynamic equilibrium and therefore the insight into prophages is of importance either on metavirome level or on level of individual isolated bacterial hosts. Several bioinformatics tools are available to detect prophages in bacterial genomes [14–18], however, these predictions still lack accuracy. Individual bacterial species of human gut commensals are understudied in terms of their prophage diversity and prevalence. Temperate phages of *Faecalibacterium prausnitzi* [19], and globally present Hankyphage, propagated from *Bacteroides dorei* were characterized and found to carry diversity generating retroelements that, through reverse transcription, introduce changes, in variable regions of genes that are often involved in host recognition [20]. Some of new *Bacteroides* phages described by our group were also detected to be integrated in bacterial genomes [21]. The recent metagenomic study of infant and parent gut metagenome samples has provided extensive insight into prophages of intestinal bacteria [11].

In this study, we aimed to screen and characterize the prophages in cultivated strains of two predominant bacterial genera in the gut microbiota, *Bacteroides* and *Phocaeicola*. In western civilizations, *Bacteroides* represent the most abundant genus in *Bacteroidota* phylum. Species of *Bacteroides* benefit to its human host, since they take part in degradation of complex polysaccharides, such as starch and xylan, produce important SCFA and contribute to colonization resistance [22]. Recently some species of *Bacteroides* genus have been transferred to newly established genus *Phocaeicola* [23]. In addition to cataloguing the prophages and their genes we also provide the insight on the stability of prophages in bacterial genomes using phage-bacterial host pairs isolated from two healthy individuals at two different time points.

## Methods

### Isolation of bacterial strains and genome recovery

Strains were selected from bacterial strain collection done by our group as described in Hedzet *et al.* [21]. In short, fecal samples, collected from two healthy donors (D1 and D2) at two temporally distant time points (S1 and S2) per each donor were stored at −80 °C. The complete isolation of bacterial strains and preparation of fecal suspension was carried out in an anaerobic workstation at 37°C. Homogenized fecal suspensions (20%) were plated on YBHI (brain–heart infusion media, supplemented with yeast extract (0.5%) and rumen fluid (20%)). After 72 h of incubation approximately 100 randomly chosen colonies per sample were subcultured on YBHI plates. Identification of pure cultures was conducted by mass spectrometry (MALDI-TOF Biotyper System, Bruker Daltonik, Bremen, Germany) and by sequencing nearly complete 16S rRNA gene. Strains identified as members of *Bacteroidaceae* (n=74) were selected for further whole genome sequencing (WGS).

Strains included in this study belong to genus *Bacteroides* (n=54), including *B. ovatus* (n=22), *B. uniformis* (n=14), *B. thetaiotaomicron* (n=14), *B. faecis* (n=4), *B. xylanisolvens* (n=3), and *B. stercoris* (n=1). Twenty isolates were from the genus *Phocaeicola*, including *P. vulgatus* (n=18) and *P. massiliensis* (n=2).

### Genome sequencing

After bacterial DNA isolation with QIAamp DNA Mini Kit (Qiagen), paired-end libraries were generated using the Nextera XT Library preparation kit (IIlumina) and sequenced on MiSeq (Ilumina) with 600-cycle MiSeq ReagentKit v3. The quality of the raw sequencing reads was examined by FastQC tool Version 0.11.9 (Babraham Bioinformatics) [24]. Quality trimming was done by Trimmomatic Version 0.39 (USADELLAB.org) [25] and overlapping paired-end reads were merged using FLASH software, version 1.2.11 [26]. Assembly was performed by SPAdes Assembler (meta spades for metaviromes), version 3.14.0 [27], and the assemblies were examined using Quast version 4.0 [28].

To identify relatedness between the obtained genomes we calculated the average nucleotide identity (ANI) using the FastANI v1.34 [29].

### Prophage analysis

To identify the most diverse collection of prophages from the obtained genomes we used PHASTER [14], Vibrant v1.2.1 [16] and Cenote-Taker v3.3.0 [18]. PHASTER was accessed as a web tool in October 2021. The quality of identified prophage regions was checked using checkV [30], and only those predicted to be complete or high-quality were kept for further analyses and termed “high-quality” prophages.

Since identified high-quality prophages are from different tools, we wanted to understand how many of them are unique. For that we clustered all sequences based on 100% similarity using MMseqs2 [31, 32]. Next, we clustered prophages into viral operational taxonomic units (vOTUs) using the MIUViG [33] recommended criteria of 95% nucleotide similarity using MMseqs2 [31, 32] with the following options: *--min-seq-id 0.95 -c 0.8 --cov-mode 0*. MMseqs2 was also used to search for the presence of obtained prophages within large viral databases. The three viral databases we used were the Metagenomic Gut Virus (MGV) catalogue with 189,680 viral genomes [34], Gut Virome Database (GVD) with 33,242 unique viral populations [35], and the Gut Phage Database (GPD) with 142,809 non-redundant gut phage genomes [36]. We created MMseqs2 databases from our own prophage sequences and sequences found in these three viral databases using the command *mmseqs created*. The search was performed with the *mmseqs search* command using the following parameters: *--max-seqs 4 --min-seq-id 0.95 --e-profile 1.0E-06 --start-sens 1 --sens-steps 3 -s 7 --threads 24 --search-type 3*.

To assign viral taxonomy to vOTUs, we used BLAST [37], vConTACT3 [38] and PhageScope [39]. This task was performed solely for representative sequences from each prophage cluster. Initially, we assigned taxonomy using vConTACT3 with the Database v220 [40] under default settings. 13 vOTUs were successfully classified at the class level. For 9 vOTUs that vConTACT3 could not classify, we employed BLAST, considering results with an E-value greater than 1e-10 and matching at least 60% of the query sequence Additionally, all sequences were processed through the PhageScope online tool using the Taxonomic Annotation option. The network results from vConTACT3 were imported into Cytoscape v3.10.2 [41] for visualization purposes.

While the phage discovery tools we used provide annotations of predicted prophage regions, they rely on different databases, leading to potential discrepancies in annotations. Therefore, we annotate all prophages with vDRAM [42].

From the same donors, we obtained fecal water metaviromes [21] to determine if prophages are potentially lytic and prevalent in these samples. For read mapping, we used coverM 0.6.1 [43] with the following parameters: *-p minimap2-sr --methods tpm -t 42 --output-format dense*.

### Data availability

Assembled bacterial genomes were retrieved from the NCBI under the accession numbers PRJNA843113, PRJNA636979. Code for read processing, prophage discovery and graphical outputs are available at: https://github.com/NLZOHomr/prophages_2024.

## Results

### Comparison of three tools for prophage detection

Altogether 597 prophages were identified in 47 out of 74 bacterial genomes, predominantly in the genus *Bacteroides* (**Fig. 1A**). Cenote-Taker 3 identified the most prophages (n=232), followed by Vibrant (n=208) and PHASTER (n=157) (**Fig. 1AB**). The quality of the identified prophages varied as well. While Cenote-Taker 3 does not provide exact quality statuses, PHASTER and Vibrant do, showing that only a small number are high-quality genomes (**Fig. 1A**). Comparing the identified prophages, we found that 21% (n=70) were detected by all three tools. Interestingly, while Vibrant and PHASTER had little overlap, Cenote-Taker 3 identified a relatively high number of prophages that overlapped with both: 67 shared with Vibrant and 25 with PHASTER (**Fig. 1B**). Furthermore, even after filtering for high-quality or complete prophages using the CheckV tool [30] the reduction in the number of prophages was significant for all three tools (**Fig. 1C**). The overlap of detected phages among three tools did not change significantly.

**Fig 1.**
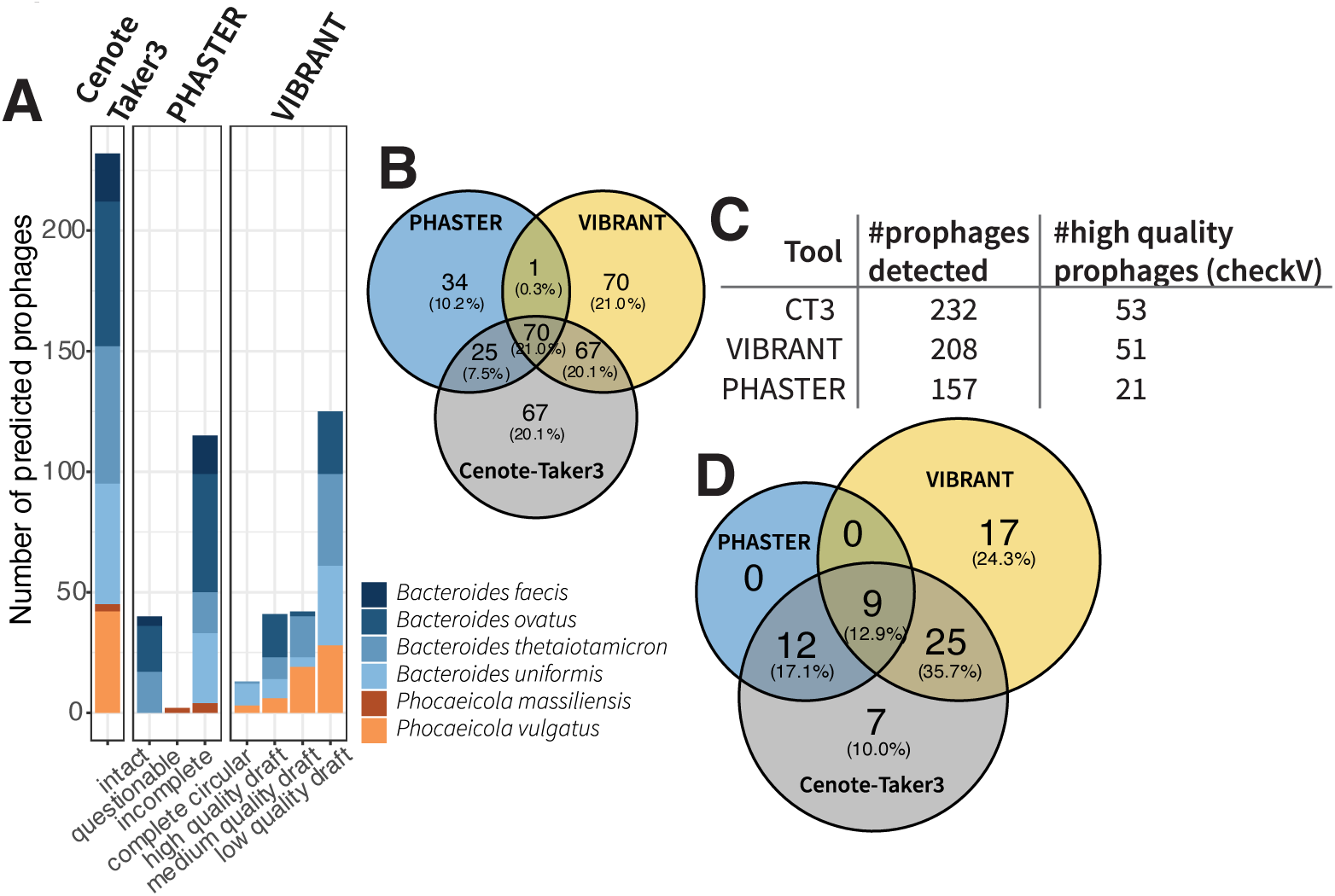
Prophage discovery. The bar plot represents number of prophages identified by the three respected tools used in the study, Cenote Taker 3, Phaster and Vibrant. These are grouped by quality as predicted by each tool. Bars are colored by the bacterial species that they were found in (A). Venn diagram highlights the number of shared and unique prophages (B). Table shows the total number of identified prophages and number after filtering for only high-quality prophages as predicted by CheckV [30] (C). Corresponding Venn diagram shows the unique and shared high-quality prophages only (D). To identify shared and unique prophages we looked at the contigs where these tools predicted prophages and if these where identical we counted as shared otherwise it was unique.

Among the tools used, Vibrant identified the most unique prophages, while Cenote-Taker 3 identified the most diverse set, covering those identified by both Vibrant and PHASTER. This makes Cenote-Taker 3 a promising sole tool for phage discovery.

### Abundance and diversity of prophages in Bacteroides and Phocaeicola strains

To identify how many unique prophages were discovered by these tools, we clustered them based on nucleotide similarity and length using MMseqs2. Out of 125 high-quality prophages, we found only 37 unique sequences at 100% similarity. At 95% similarity, 22 clusters were formed, which we refer to as viral operational taxonomic units (vOTUs) (**Table 1**). vOTU 4 was the only cluster containing prophages identified by all three tools (**Table 1**). The size of the identified prophages ranged from 27,455 bp (vOTU11) to 124,118 bp (vOTU19), encoding between 32 and 164 genes.

**Table 1.** Summary of the vOTUs. Prophages that were clustered at 95% similarity formed 22 distinct vOTUs. Prophages that are within these vOTUs were identified by either one, two or once by all three tools (V – Vibrant, C – Cenote Taker 3, P – Phaster). Length of prophage genomes within each cluster and the number ORFs are also represented. Information in the brackets represents minimum and maximum values within each cluster. Taxonomy column contains viral class as predicted by either vContACT3, BLAST or PhageScope.

| vOTU | Tools | Length (bp) | ORF | Taxonomy |
| --- | --- | --- | --- | --- |
| vOTU1 | V, C | 42160.88 (37638 – 48793) | 51.5 (45 – 63) | *Caudoviricetes* |
| vOTU2 | C, P | 42759.29 (42,608 – 42961) | 45 | *Caudoviricetes* |
| vOTU3 | C | 35454 | 38 | *Caudoviricetes* |
| vOTU4 | V, C, P | 50266 (49032 – 51774) | 63.3 (62 – 65) | *Caudoviricetes* |
| vOTU5 | C | 29500 | 38 | *Caudoviricetes* |
| vOTU6 | C | 61508 (59309 – 63707) | 74 (72 – 76) | *Caudoviricetes* |
| vOTU7 | C | 43600 | 54 | *Caudoviricetes* |
| vOTU8 | C | 63319 | 82 | *Caudoviricetes* |
| vOTU9 | V, C | 73695 (71863 – 75527) | 109 (108 – 110) | *Caudoviricetes* |
| vOTU10 | V, C | 43789.11 (40315 – 45709) | 46.9 (41 – 49) | *Caudoviricetes* |
| vOTU11 | P | 27455 | 32 | *Caudoviricetes* |
| vOTU12 | P | 33914 | 36 | *Caudoviricetes* |
| vOTU13 | V, C | 75645.25 (71365 – 81021) | 109 (103 – 116) | *Caudoviricetes* |
| vOTU14 | V, C | 48011 (44905 – 51117) | 56 (52 – 60) | *Caudoviricetes* |
| vOTU15 | V, C | 52719.75 (50763 – 53372) | 66.8 (66 – 67) | *Caudoviricetes* |
| vOTU16 | V | 81,403 | 116 | *Caudoviricetes* |
| vOTU17 | V | 63326 (63296 – 63,356) | 72 | *Caudoviricetes* |
| vOTU18 | V | 62,533 | 72 | *Caudoviricetes* |
| vOTU19 | V | 120873.50 (111182 – 124118) | 161.3 (153 – 164) | *Caudoviricetes* |
| vOTU20 | V, C | 88,336 | 110 | *Caudoviricetes* |
| vOTU21 | C | 113780 (112145 – 115415) | 155 (154 – 156) | *Caudoviricetes* |
| vOTU22 | V, C | 69564 (67265 – 71863) | 100 (93 – 107) | *Caudoviricetes* |

**Table 1.**
| vOTU | Tools | Length (bp) | ORF | Taxonomy |
| --- | --- | --- | --- | --- |
| vOTU1 | V, C | 42160.88 (37638 – 48793) | 51.5 (45 – 63) | Caudoviricetes |
| vOTU2 | C, P | 42759.29 (42,608 – 42961) | 45 | Caudoviricetes |
| vOTU3 | C | 35454 | 38 | Caudoviricetes |
| vOTU4 | V, C, P | 50266 (49032 – 51774) | 63.3 (62 – 65) | Caudoviricetes |
| vOTU5 | C | 29500 | 38 | Caudoviricetes |
| vOTU6 | C | 61508 (59309 – 63707) | 74 (72 – 76) | Caudoviricetes |
| vOTU7 | C | 43600 | 54 | Caudoviricetes |
| vOTU8 | C | 63319 | 82 | Caudoviricetes |
| vOTU9 | V, C | 73695 (71863 – 75527) | 109 (108 – 110) | Caudoviricetes |
| vOTU10 | V, C | 43789.11 (40315 – 45709) | 46.9 (41 – 49) | Caudoviricetes |
| vOTU11 | P | 27455 | 32 | Caudoviricetes |
| vOTU12 | P | 33914 | 36 | Caudoviricetes |
| vOTU13 | V, C | 75645.25 (71365 – 81021) | 109 (103 – 116) | Caudoviricetes |
| vOTU14 | V, C | 48011 (44905 – 51117) | 56 (52 – 60) | Caudoviricetes |
| vOTU15 | V, C | 52719.75 (50763 – 53372) | 66.8 (66 – 67) | Caudoviricetes |
| vOTU16 | V | 81,403 | 116 | Caudoviricetes |
| vOTU17 | V | 63326 (63296 – 63,356) | 72 | Caudoviricetes |
| vOTU18 | V | 62,533 | 72 | Caudoviricetes |
| vOTU19 | V | 120873.50 (111182 – 124118) | 161.3 (153 – 164) | Caudoviricetes |
| vOTU20 | V, C | 88,336 | 110 | Caudoviricetes |
| vOTU21 | C | 113780 (112145 – 115415) | 155 (154 – 156) | Caudoviricetes |
| vOTU22 | V, C | 69564 (67265 – 71863) | 100 (93 – 107) | Caudoviricetes |

Per genome, we identified between 1 and 5 high-quality vOTUs, with the highest number of prophages found in two strains of *P. vulgatus* (MB20-93 and MB20-96) (**Fig. 2**). While bacterial genomes from a particular species show high ANI (**Fig. 2A, Suppl Fig. S1**), the number of high-quality prophages can vary significantly. This variation may be due to unequal integration of prophages within the population or missing genome information, as many genomes are in draft stages (56–392 contigs per genome). vOTUs 4, 10, and 16 were found in the highest number of bacterial genomes (n=9) (**Fig. 2B**).

**Fig 2.**
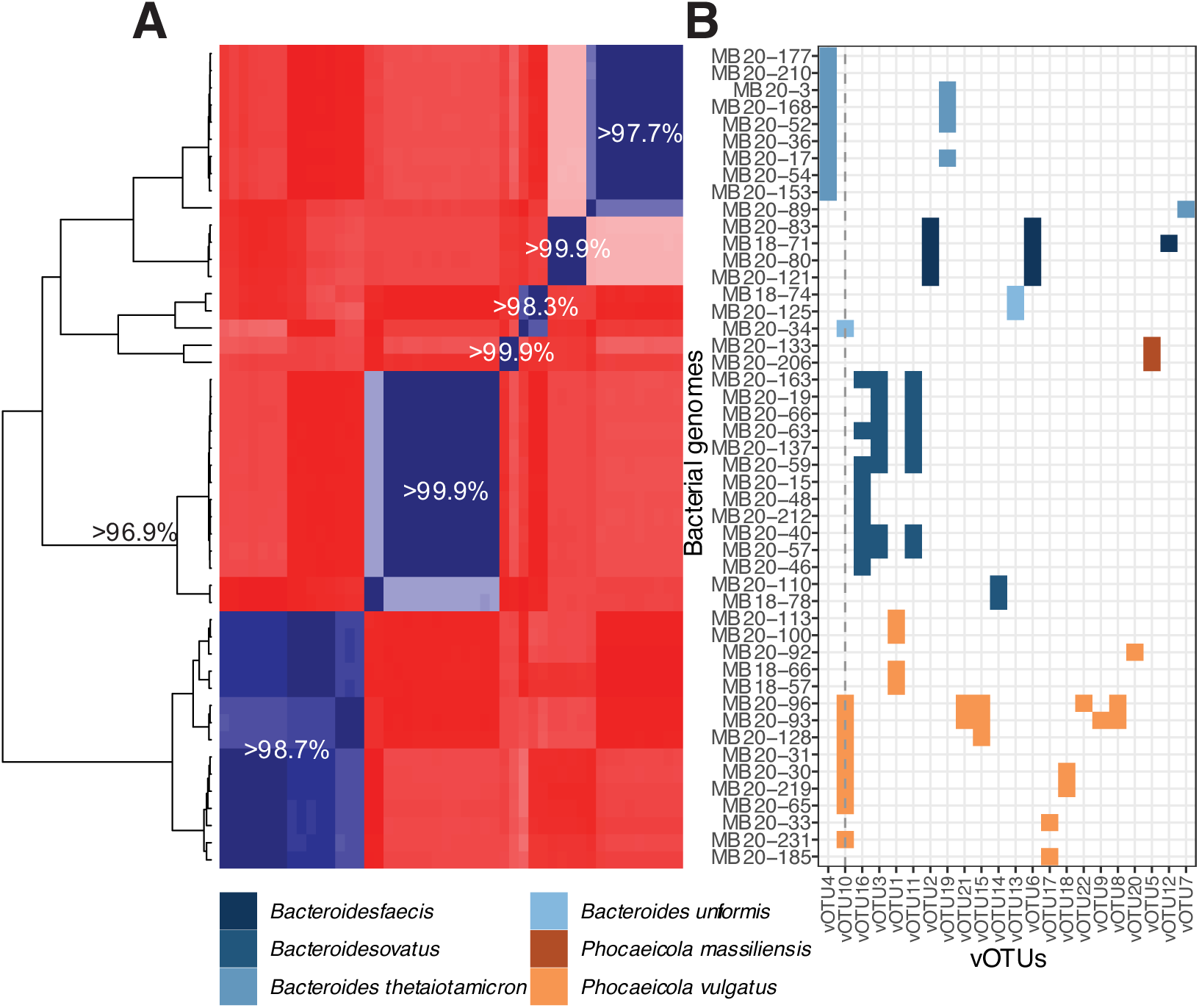
vOTU host-range. Bacterial genomes where high-quality genomes were identified are represented in the ANI heatmap. The minimum ANI scores for each block are highlighted on the heatmap and are above 96.9%. Corresponding bacterial IDs are marked on right side of the heatmap. vOTUs per bacterial genome are represented as dot plot where colors represent bacterial species where vOTUs were found. vOTU10 is highlighted by break line as the only vOTU which is found in two different host species.

**Fig S1.**
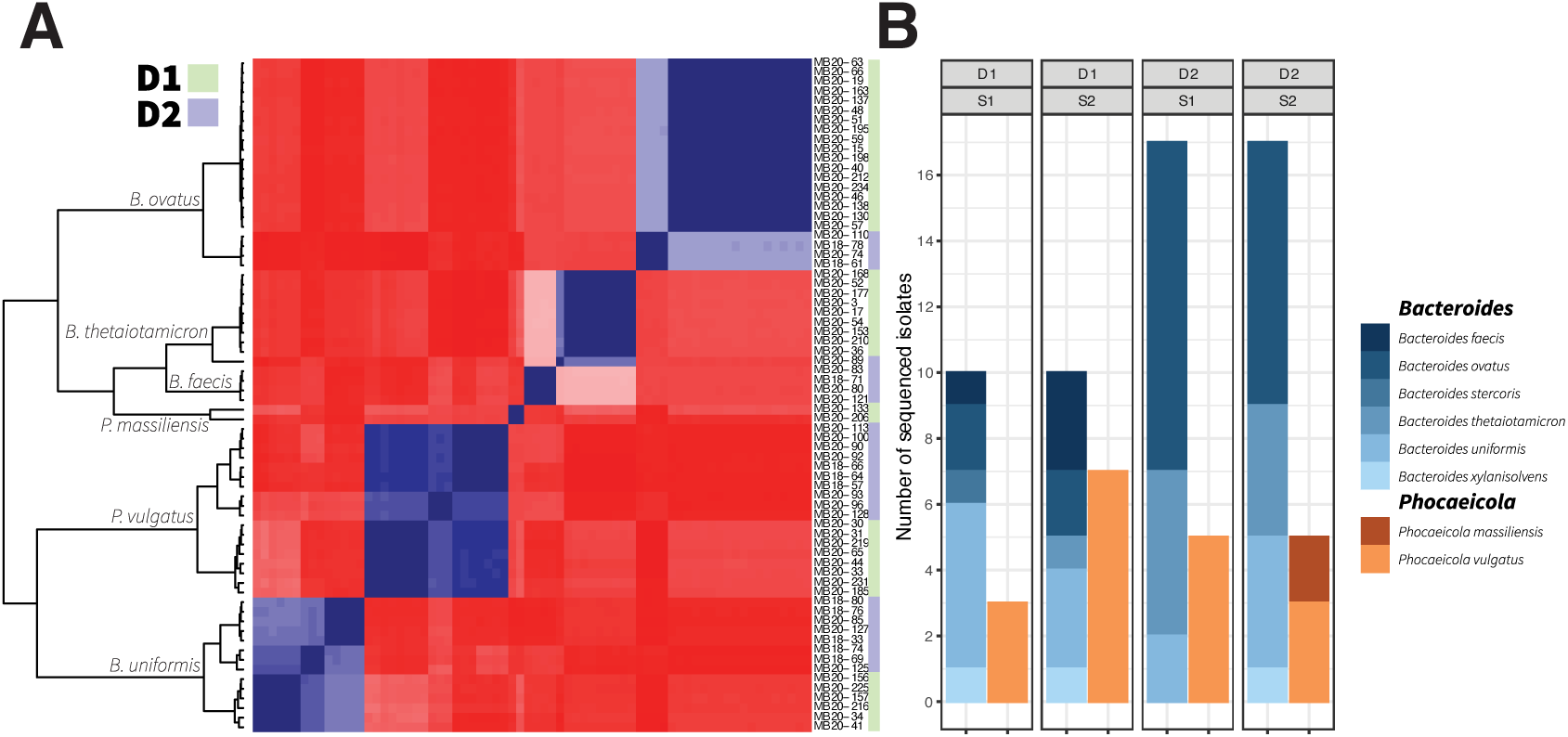
Bacterial genome summary. The ANI scores are represented as heatmap where dark blue colors represent 100% identity and dark red identity bellow 77% as calculated by FastANI [29] (**A**). Bacterial species are marked on the branches of the dendrogram and donors where the bacterial cultures derived from are highlighted by color on the right side of the heatmap next to the bacterial IDs (green – donor 1 (D1) and purple – donor 2 (D2)). Number of bacterial genomes by donor, time point and phylogenetic association are represented in the bar plot (**B**).

Furthermore, vOTU10 was the only vOTU identified in two different genera, *B. uniformis* and *P. vulgatus*, while all other vOTUs were highly species-specific and probably also strain-specific (**Fig. 2**).

Taxonomic classification revealed that all identified prophage clusters belong to the highly diverse tailed bacteriophage class *Caudoviricetes*. vConTACT3 predicted 13 vOTUs to this class and 9 were unassigned. However, using BLAST and PhageScope we were able to predict taxonomy also of the remaining 9 vOTUs which all belong to *Caudoviricetes*. Based on vConTACT3 vOTUs make up three larger clusters, comprising of 10, 5 and 2 vOTUs, respectively and 4 vOTUs which do not have any connectivity (**Fig. S2**).

**Fig S2.**
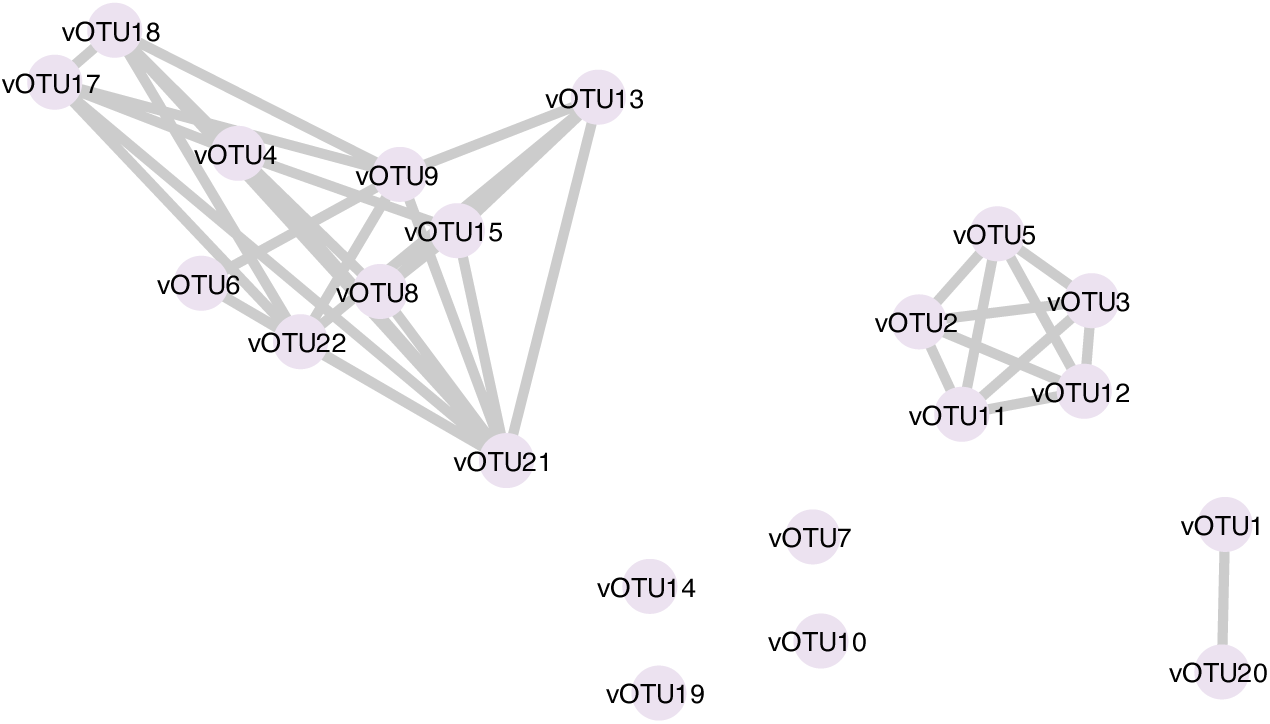
Taxonomic analysis of vOTUs. We used vConTACT3 [38] for inferring the taxonomy of viral sequences which is based on the shared protein clusters. The network was visualized and modified in Cytoscape v3.10.2 [41].

### Prophage temporal dynamics in the gut environment of our donors

Our study allowed us to investigate not only the host range of these prophages but also their dynamics over two time periods. We found that vOTUs seemed host-specific and also highly donor-specific (**Fig. 3**). Only vOTU10 was found in both donors, while all others were unique to a single donor. Specifically, vOTU10 was shared between two genera only in donor 2 at sampling time 1 (**Fig. 3**). Additionally, when comparing the occurrences of vOTUs over time, we observed that vOTUs are temporally stable (**Fig. 3**). Except for vOTU12, any vOTU present at sampling time 1 (S1) also appeared at sampling time 2 (S2) within the same donor (**Fig. 3**). However, prophages found at sampling time 2 were not always present at sampling time 1, likely due to uneven bacterial culturing across different sampling periods.

**Fig 3.**
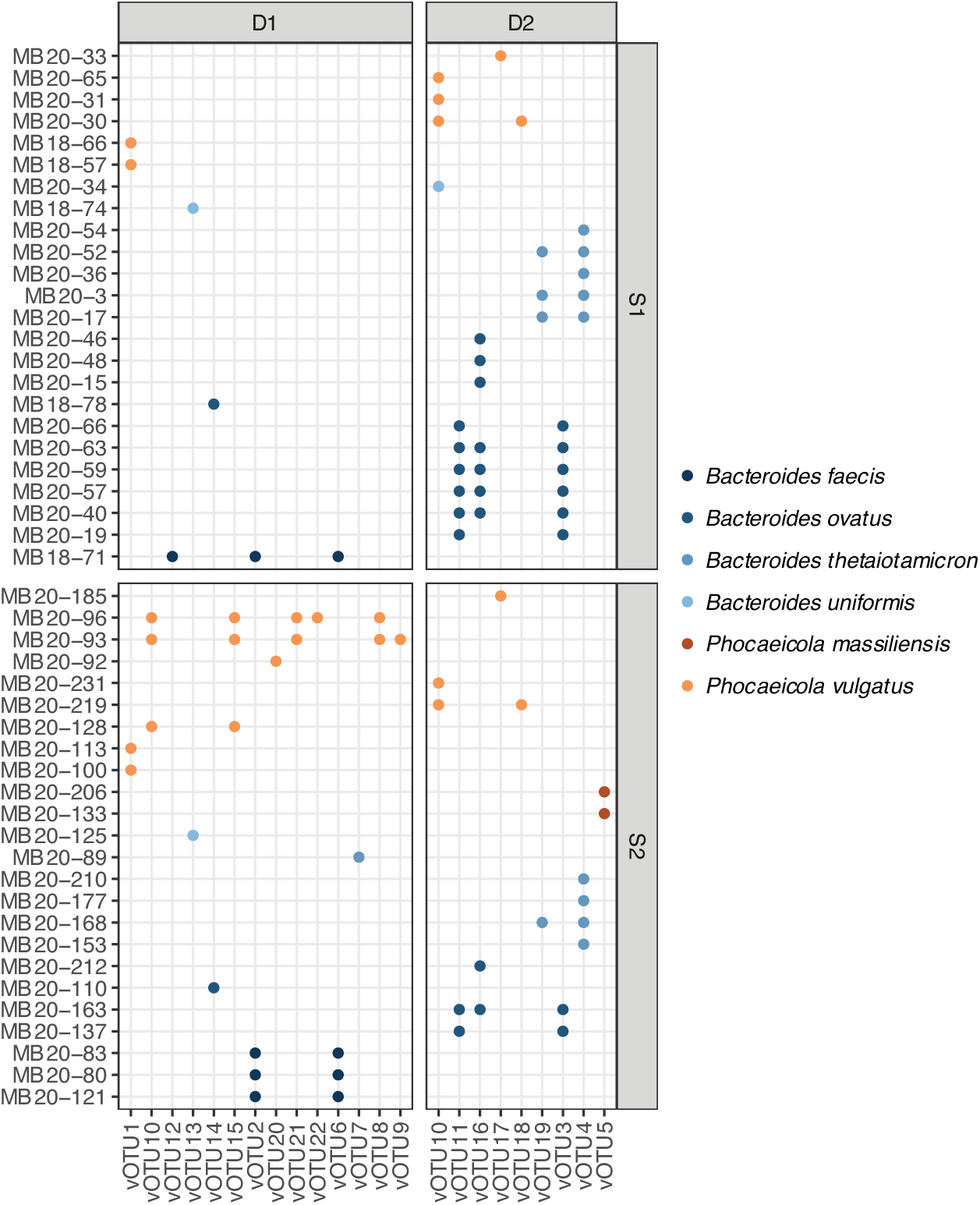
vOTU over donor and time. The dot plot represents the occurrence of vOTUs in donor 1 (D1) or donor 2 (D2) at time 1 (S1) or time 2 (S2). Each dot is color coded by bacterial host.

### Prophage host range at the level of bacterial isolates, metaviromes and global gut databases

Previously [21], we obtained three metavirome samples from the same individuals, which we used for mapping onto the reference vOTUs. Reads from all metaviromes mapped to the vOTUs, indicating that these prophages are not only lysogenic but can also exhibit a lytic cycle. Surprisingly, these prophages can become relatively abundant, as suggested by their high mapping percentages. The sample from individual 1 (D1_FW2) had the highest mapping percentage, accounting for 6% of the total metavirome, while reads from the individual 2 (D2) mapped at 1.35% and 1.17%, for time points 1 and 2, respectively. Interestingly, reads mapped to 20 out of 22 prophage vOTUs, mainly across both donors (**Fig. 4**), suggesting that these prophages are prevalent in the human gut of these donors.

**Fig 4.**
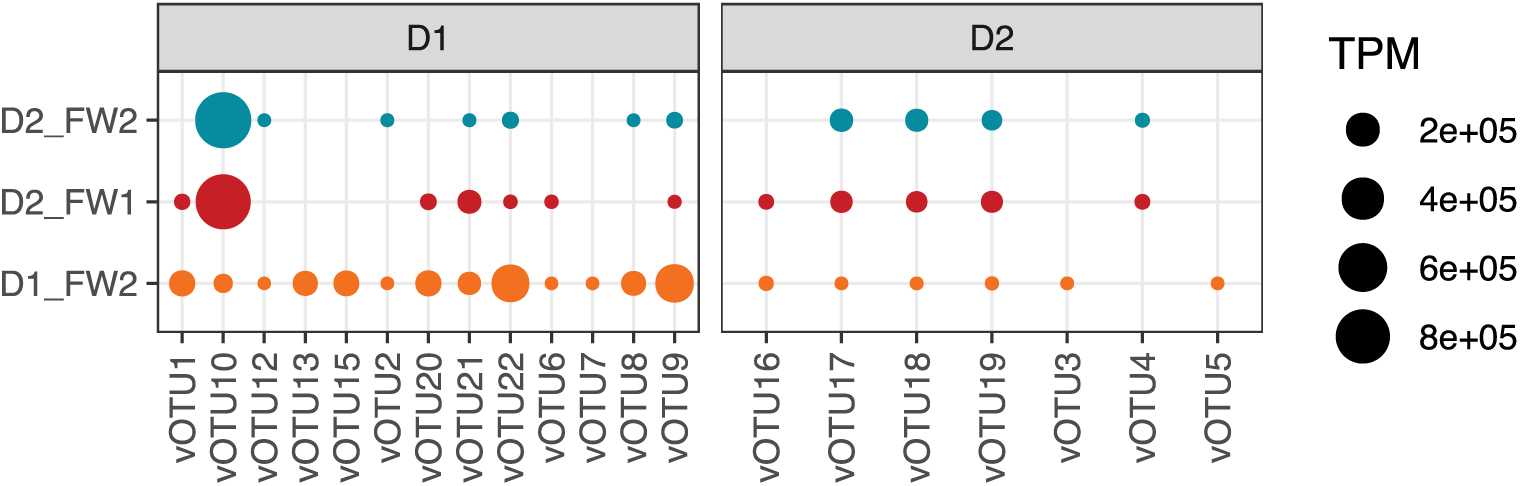
Mapping summary of metavirome samples (fecal water). We mapped short reads on database of unique high-quality prophages (n=37) using coverM [43] and expressed the coverage measurement as transcripts per million (TPM) which are represented as dot sizes. Dots are colored by metavirome samples.

These results raised the question of whether vOTUs are as host- and donor-specific as our bacterial genome analysis suggests. To explore this, we examined three publicly available gut virome databases. All vOTUs identified in the present study were found in all three databases (**Supplemental Table S1**). While the genus *Bacteroides* is predicted to be the sole host for all our vOTUs in the MGV database, the GPD and GVD databases predict additional hosts. For instance, vOTU5 was previously found in members of the family *Tannerellaceae*, vOTU10 in *Bacteroides dorei* and the family *Enterococcaceae*, and vOTU19 in *Bacteroides faecis*. Moreover, the GPD database includes geographical locations of the best-matched hits, showing that the studies where our prophages were discovered are spread across the globe (e.g., China, United States, Denmark, Germany, Australia, Russia, etc.) (**Supplemental Table S1**).

### Prophage genetic content

To understand the genetic content of the identified prophages, we annotated reference vOTU genomes from 22 clusters using the vDRAM tool [42]. The gene catalogue comprises 1,659 open reading frames (ORFs), with approximately 43% of these ORFs successfully annotated (**see Supplemental Table S2**).

Fourteen vOTUs were predicted to carry auxiliary metabolic genes (AMGs) involved in various metabolic processes (**Supplemental Table S2**). Among the identified AMGs, ketopantoate hydroxymethyltransferase, an enzyme essential for vitamin B5 (pantothenate) biosynthesis, was detected. Several sugar transferases, such as glycosyl transferase, were found, which could enhance carbon metabolism and cell wall biosynthesis in the host bacteria. Notably, vOTU21 encodes a potential pullulanase, an enzyme involved in the breakdown of starches and polysaccharides, while vOTU1 encodes alpha-L-rhamnosidase, which is involved in the degradation of plant materials and microbial polysaccharides.

Several toxin genes were identified in many vOTUs, indicating potential roles in bacterial competition and survival (**Supplemental Table S2**). Thiol-activated cytolysin [44] was identified in vOTU8, while toxin SymE [45], part of a type I toxin-antitoxin system, was found in vOTU14. Additionally, a putative AbiEii toxin from a type IV toxin-antitoxin system [46] was present in vOTU14, and the antitoxin component MqsA was found in vOTUs 8 and 15. A type II toxin-antitoxin system HigB family toxin, with an associated antitoxin HigA, was identified in vOTUs 16 and 21 [47].

Furthermore, several vOTUs contained genes associated with antibiotic resistance (**Supplemental Table S2**). Chloramphenicol O-acetyltransferase type A, which detoxifies chloramphenicol and confers resistance, was identified in vOTUs 2 and 3. Multidrug resistance proteins of the MATE family were found in vOTUs 2 and 12. Beta-lactam resistance genes, including those encoding beta-lactamase superfamily domains, were found in vOTUs 4, 8, and 15, while L,D-transpeptidase catalytic domains were present in vOTU16.

## Discussion

In this study, we investigated the diversity, host range, and temporal dynamics of prophages within the *Bacteroidaceae* family using a combination of culture-based methods and comprehensive genomic analyses.

The use of multiple bioinformatic tools - PHASTER, Vibrant, and Cenote-Taker 3 - proved crucial in capturing the full spectrum of prophage diversity. Each tool employs different algorithms and reference databases, resulting in varied prophage predictions [16]. The complementary nature of these tools allowed us to identify a broader range of prophages than any single tool could achieve alone. This approach aligns with findings from other studies, which emphasize the necessity of using diverse methodologies to obtain a comprehensive view of viral diversity [10, 48, 49].

Culture-based methods, while invaluable, often present a skewed view of microbial diversity due to their selective nature. This study mitigated such biases by integrating metavirome data and leveraging large viral databases, providing a holistic perspective on prophage diversity and host range. Our initial findings from cultured bacterial strains indicated a narrow host specificity for many prophages, confined to specific species or even strains. However, metavirome analysis and database comparisons broadened this perspective, revealing that these prophages also infect a wider array of bacterial hosts. This integrative approach is crucial for understanding the true ecological roles of prophages.

All identified prophages belong to the viral class *Caudoviricetes*, a diverse group of tailed bacteriophages that are highly prevalent in the gut microbiome [50, 51]. This class is known for its role in horizontal gene transfer, which can significantly influence bacterial evolution and adaptation [52]. The presence of *Caudoviricetes* prophages in the gut has been linked to various beneficial and detrimental effects on the human host, including modulation of immune responses and contribution to gut dysbiosis [53, 54]. Our findings contribute to the growing body of evidence that *Caudoviricetes* prophages are integral components of the gut microbiome, capable of shaping bacterial community structure and function.

In terms of host range and dynamics, vOTU10 stood out among the clusters examined, being consistently present in both donors across both time points. Our culture data initially identified vOTU10 in *B. uniformis* and *P. vulgatus*. Subsequent analysis using viral databases expanded its host range to include *B. dorei* and *Enterococcaceae*. BLAST results indicated a close match (99.10% identity and 94% query coverage) with members of the *Bacteroides* phage Hanky p00 (Hankyphage) group, known as temperate phages of *B. dorei* capable of lysing various *Bacteroides* species and prevalent in metaviromes [55]. Importantly, this prophage, along with vOTU1 and vOTU20, was previously identified in the same samples through meticulous manual curation of PHASTER predictions, despite these prophages not being initially predicted as high quality [21, 56]. Our integrated analysis approach now allows us to efficiently resolve these prophages using high-throughput bioinformatics tools, providing confidence in our findings and demonstrating that we can reliably detect the same prophages without the need for labor-intensive manual curation. These findings underscore the effectiveness of combining multiple phage discovery tools to ensure comprehensive and accurate phage identification.

One of the important findings in our study is the presence of several antibiotic resistance genes (AMR) and genes that could potentially enhance and contribute to host metabolism within the prophages. This is consistent with previous research, which has documented the presence of AMR genes and auxiliary metabolic genes in prophages, emphasizing their role in bacterial adaptability and survival [57–60]. Our identification of AMR genes aligns with findings from other studies that have reported similar genes in gut-associated phages [61]. The presence of these genes in prophages is of significant concern as they can facilitate horizontal gene transfer, potentially spreading resistance traits among bacterial populations in the gut microbiome. This contributes to the growing problem of antibiotic resistance, which is a critical public health issue.

The presence of toxin genes, including thiol-activated cytolysin and various toxin-antitoxin systems, indicates potential roles in bacterial competition and survival. These toxins can help prophage-containing bacteria outcompete other microbial populations, thereby influencing the composition and dynamics of the gut microbiome [9, 44, 62, 63].

In addition to AMR and virulence genes, we discovered genes encoding proteins involved in various metabolic processes. These AMGs can enhance the metabolic capabilities of their bacterial hosts, supporting functions like vitamin B5 biosynthesis, carbon metabolism, and the degradation of complex polysaccharides [63–65].

In summary, we have used integrated approach for prophage analysis which included three different tools for phage detection, comparison of results between bacterial genomes and metaviromes and annotation of prophage genomic analysis. Our results show that identified prophages are widely found in the human gut ecosystem and are likely able infecting abroad range of bacterial species. This promiscuity may explain their stable presence over periods ranging from three months to a year. Genomic analysis revealed a variety of important functional genes including those involved in specific metabolic pathways, antibiotic resistances and toxin production.

## Declarations

### Ethics approval and consent to participate

This work utilized sequencing data from a previous study, for which written informed consent was obtained from all participants. Ethical approval was granted by the National Medical Ethics Committee of the Republic of Slovenia (reference number: 0120-142/2016-2).

### Consent for publication

Not applicable.

---

**Supplemental Table S1.** Summary of the vOTU search against the large gut virome databases. The column names represent the following:

- *vOTU* = vOTU ID
- *DB* = reference database (see Methods for details)
- *DB Reference* = source of the studz where vOTU was identified
- *Host* = bacterial hosts as assigned from the databases
- *Genome size* = genome size of the matched reference
- *Different host range from our study* = with Y we highlighted those vOTUs for which bacterial host is different from ours.
- *Study countries* = countries where vOTUs have a match (available only for GPD database)

**Supplemental Table S2**. Combined result output from annotation with vDRAM.

